## Supplementary Information for "Raman micro-spectroscopy reveals the spatial distribution of fumarate in cells and tissues"

**Title**

### Supplementary Note 1

Given the importance of the mitochondrial peak assignment in the k-means clustering for evaluation of fumarate concentrations, we used mitokyne (main text reference 42), a recently introduced Raman probe that tags mitochondria via a triphenylphosphonium group to enable detection of mitochondria by the appearance of a vibrational mode in the silent region. The primary mitokyne band (located close to 2217 $cm^{-1}$, pH-dependent) and secondary band (at 1613 $cm^{-1}$) are both visible in the averaged area scans of tagged WT or KO cells (Figure S14A). No mitokyne bands occlude the fumarate 1401 $cm^{-1}$ mode. A k-means clustering was performed jointly over the area scans for Fh1-KO cells without (20 cells) and with (10 cells) mitokyne, yielding a new set of cluster maps and loadings, *K2*. K-means cluster maps K2 were identical to K1 for the non-labelled Fh1-KO cells (c.f. Figure 3E and S14B, right). Loadings of K1,K2 were similar to a high degree with Spearman correlation coefficients close to 1 for the loadings of the nucleus (0.999945), membrane (0.999934), cytoplasm (0.999945) and mitochondria (0.999880), indicating the loadings rise and fall at the same wavenumbers with the lower correlation for the mitochondria stemming from the mitokyne bands. The mitokyne primary band was indeed most intense in the loadings corresponding the mitochondria (Figure S14B), evidencing that our k-means cluster assignment for the mitochondria is correct. Further, a positive correlation was observed across all cell compartments between the intensity of the mitokyne primary band and the fumarate 1401 $cm^{-1}$ band (Figure S14C-D), suggesting a preferential localization of fumarate in the mitochondria, as would be expected.

<u>**Supplementary Figures**</u>

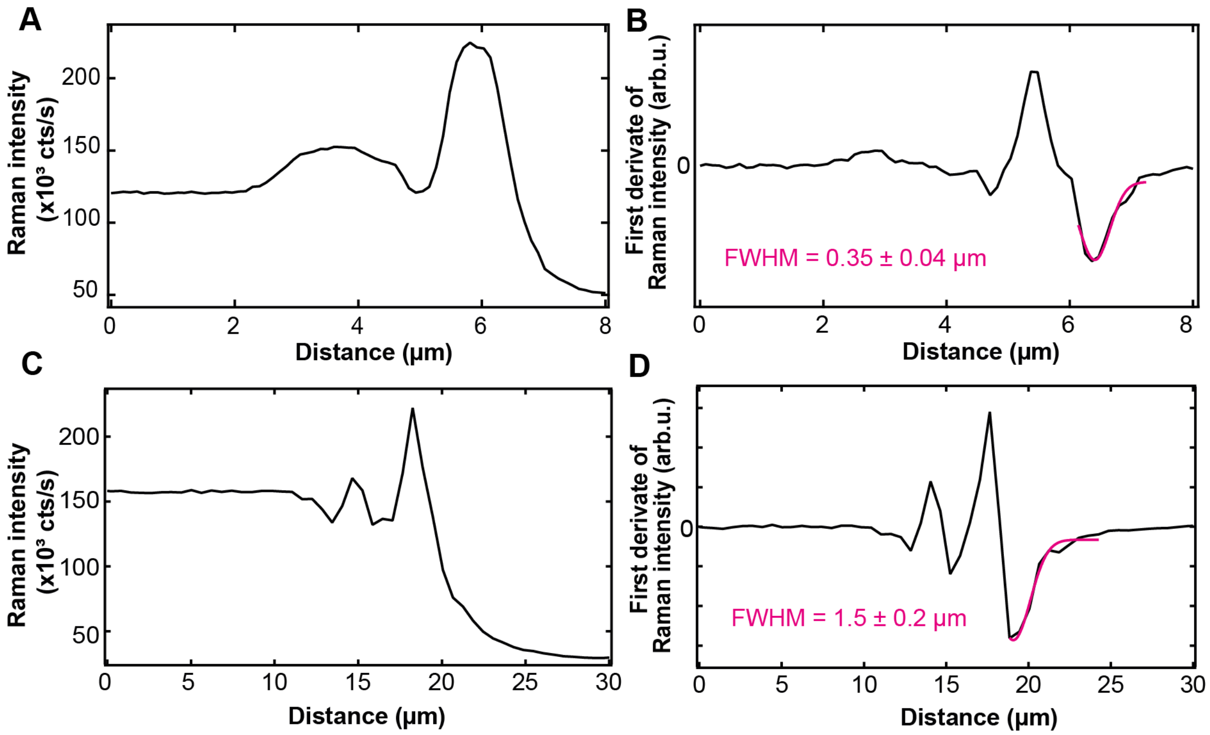


**Figure S1. Confirmation of instrument spatial resolution.** (**A**) Raman intensity of the silicon peak along a line over the edge of the silicon wafer (edge response). $\lambda_{exc}$ = 532 nm. (**B**) Derivative of the edge response (LSF). **C**) Raman intensity of the silicon peak along a line over the edge of the silicon wafer (edge response). $\lambda_{exc}$ = 785 nm. **D**) Derivative of the edge response (LSF).

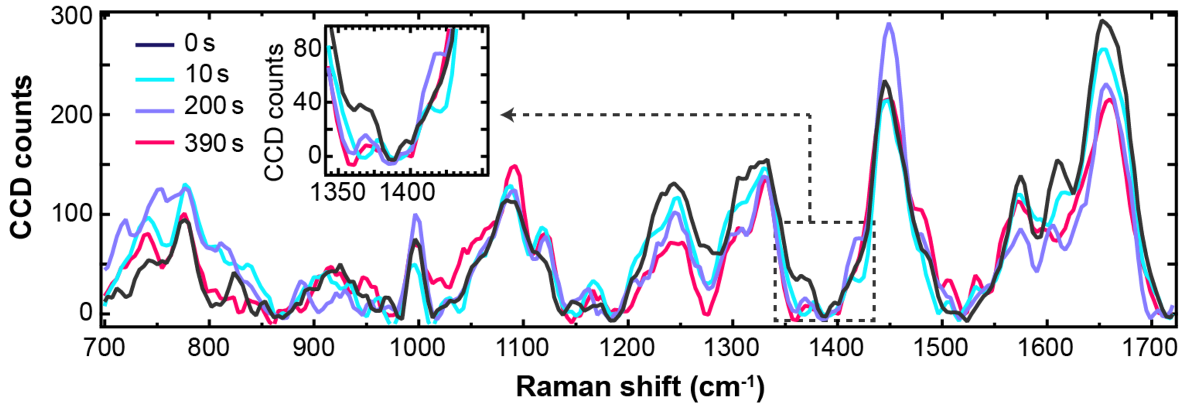


**Figure S2. Cell damage due to laser illumination.** A single point on an Fh1-WT cell is illuminated ($\lambda_{exc}$ = 532 nm) 400 times (0.5 s on and 0.5 s off). The spectra shown are those at time points 0 s, 10 s, 200 s, and 390 s.

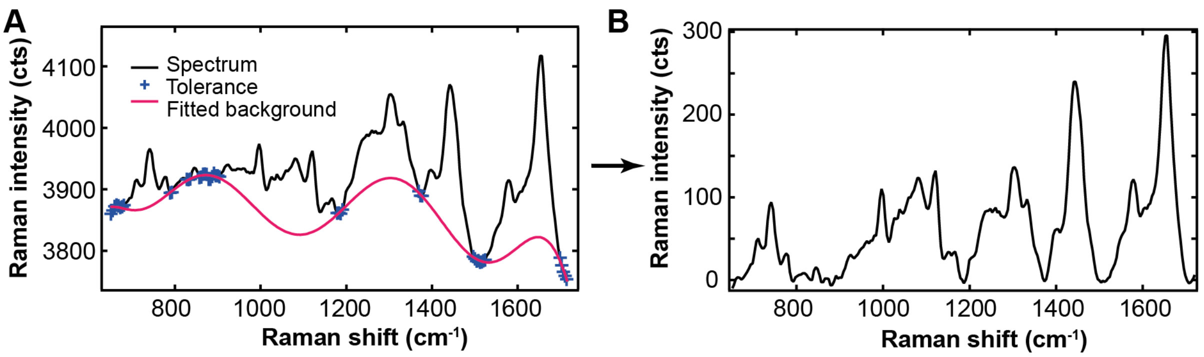


**Figure S3. Visualization of the iterative baseline subtraction used to compare average cell spectra.** (**A**) A 10th order polynomial was fitted to the entire spectrum (black), after which all graph points over a chosen 'tolerance' above the fitted polynomial (in counts, in the current work: 6 cts) are removed from the spectrum to be used for the subsequent fit. This procedure is iterated ten times. The blue crosses indicate the remaining data points through which the tenth (final) polynomial fit was conducted. The pink line is the resulting baseline fit. (**B**) Baseline corrected spectrum.

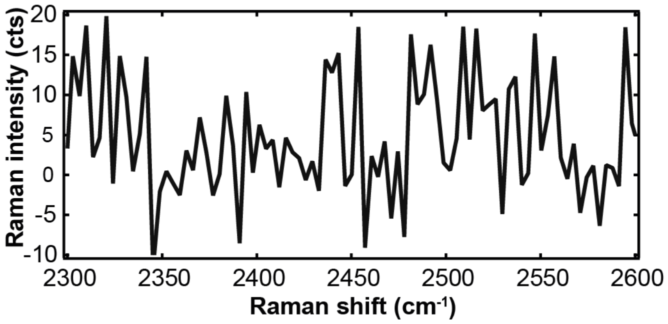


**Figure S4. Instrument noise test.** A typical spectrum was baseline-corrected in the silent region of the wavenumber range. Over 100 pixels, the standard deviation of the Raman intensity was 7 cts.

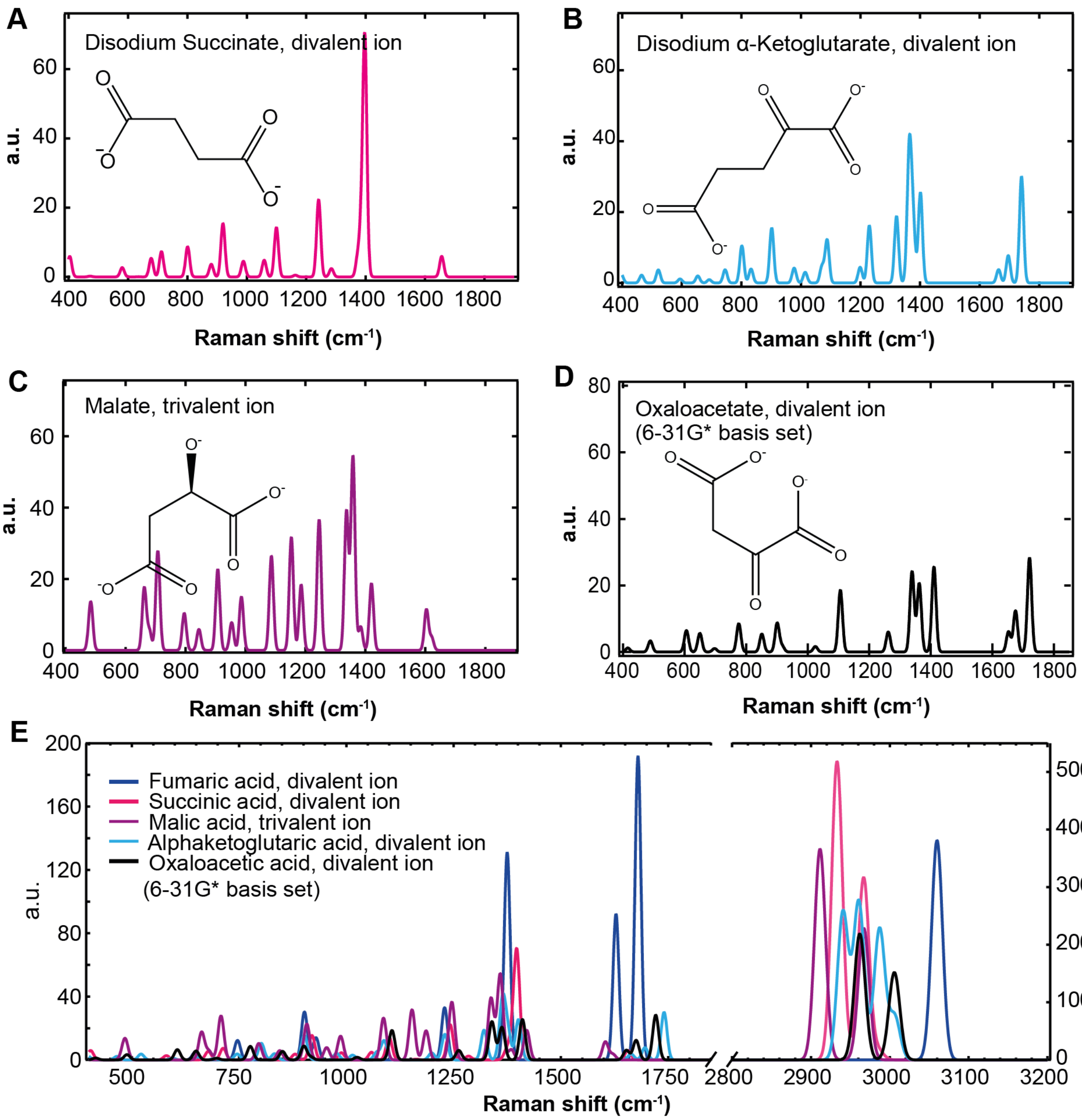


**Figure S5. DFT Calculations of the Raman spectra for major metabolites of the TCA cycle.** Divalent ions of: (**A**) succinate, (**B**) α-ketoglutarate, (**C**) malate, (**D**) oxaloacetate. (**E**) DFT-calculated Raman spectra of the same metabolites and of fumarate including the long-wavenumber regime. The divalent fumarate ion can be seen to have a >2x higher peak intensity in the fingerprint region than other metabolites.

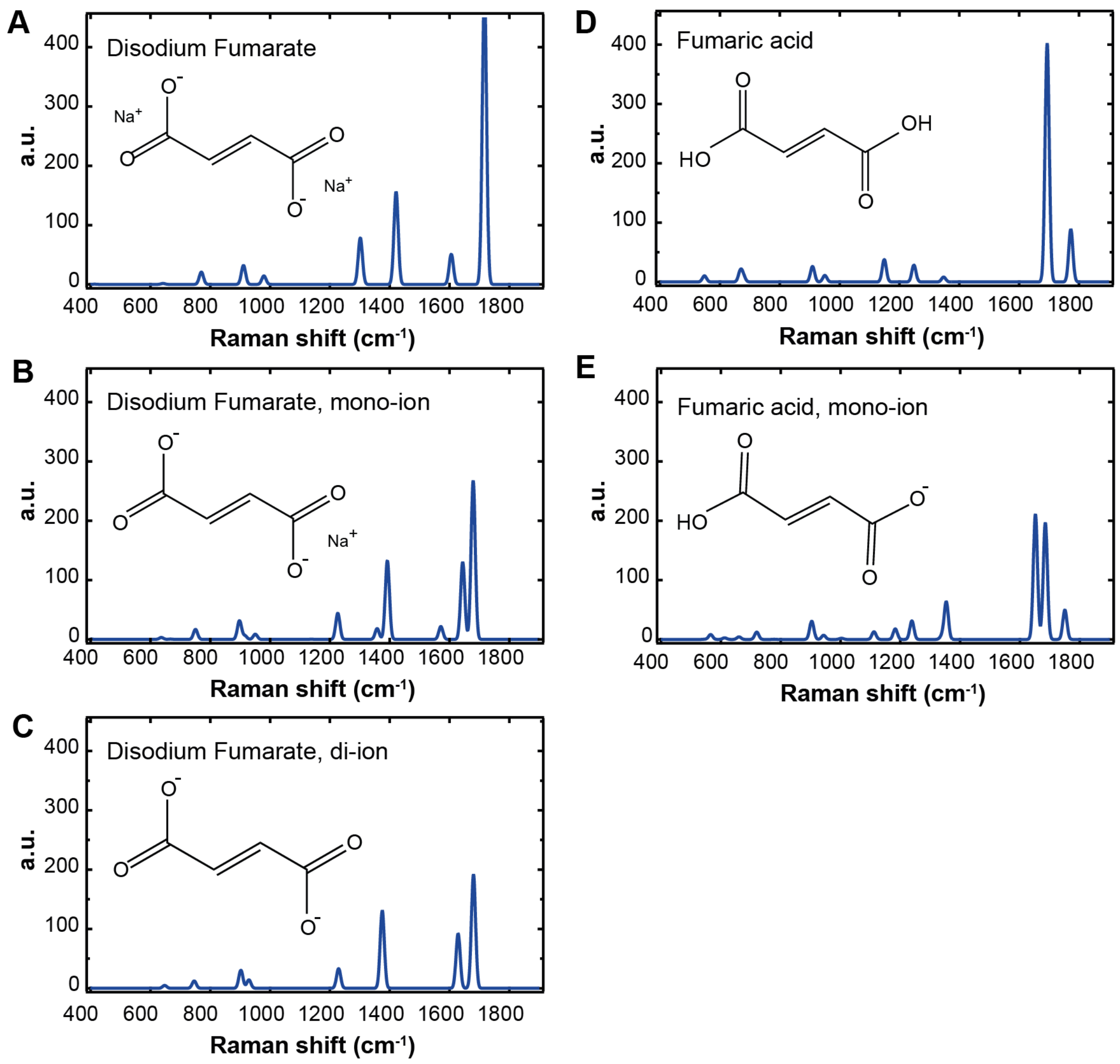


**Figure S6. DFT Calculations of the Raman spectra of fumarate molecules.** Raman spectra obtained via DFT calculations for fumarate in different ionization states: (**A**) disodium fumarate, (**B**) monovalent ion of disodium fumarate, (**C**) divalent fumarate ion, (**D**) fumaric acid, and (**E**) monovalent ion of fumaric acid.

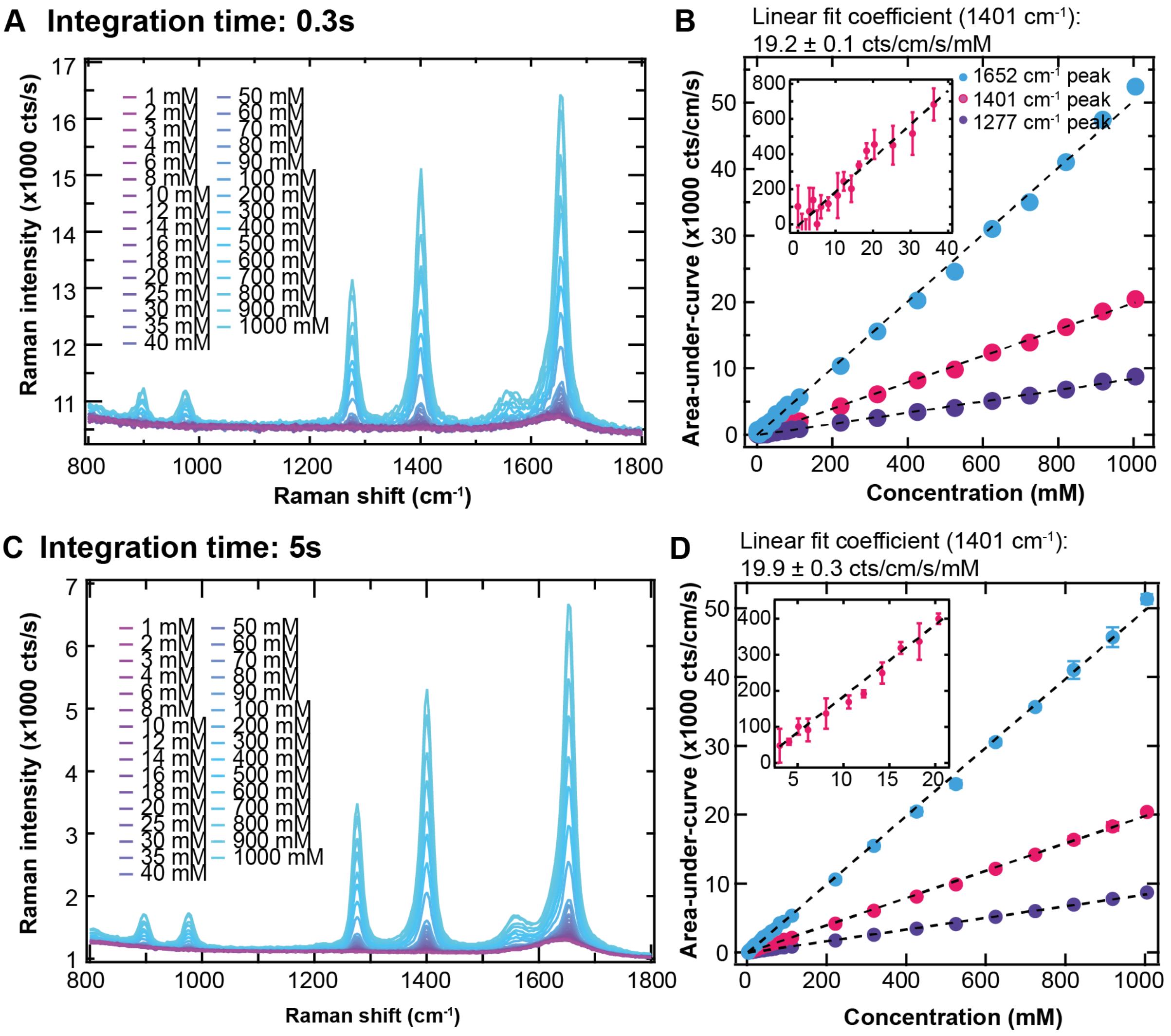


**Figure S7. Concentration dilution series of aqueous disodium fumarate showing the influence of dwell time**, analysed by the area under the curve. Raman intensities in counts/s at 26 mW laser power, 532 nm laser excitation for dwell times of 0.3s (**A**) and associated linear fit to dilution series (**B**). (**C,D**) Equivalent for 5 s dwell time. The error bars represent the standard deviation of three different measurements obtained at the same concentration. The $r^2$ values for the linear regression are in (b): 0.998791 (1652 $cm^{-1}$), 0.998556 (1401 $cm^{-1}$) and 0.997838 (1277 $cm^{-1}$); and in (d): 0.999136 (1652 $cm^{-1}$), 0.997005 (1401 $cm^{-1}$), and 0.998963 (1277 $cm^{-1}$).

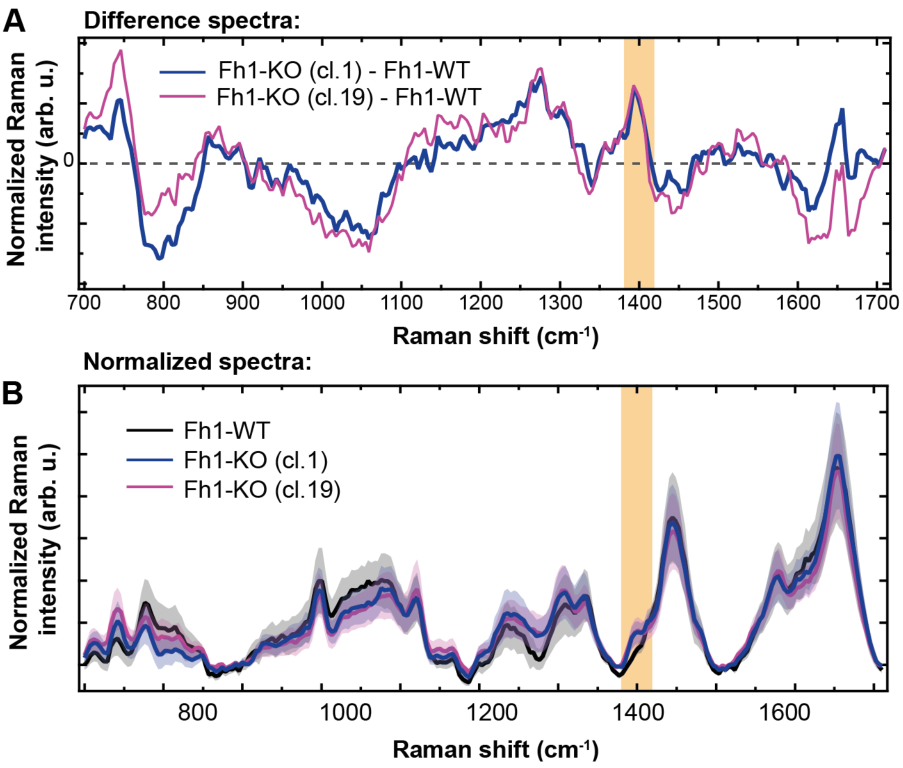


**Figure S8. Investigation of fumarate peak visibility using different post-processing**. (**A**) Difference spectra emphasizing the fumarate peak at 1401 $cm^{-1}$: Averaged spectra of 20 Fh1-KO cl.1 cells (purple) and 20 Fh1-KO cl.19 (pink) minus the averaged spectra of 20 Fh1-WT cells. (**B**) Averaged area scans normalized by the area under the curve in the wavenumber range 650-1720 $cm^{-1}$. Normalization equalizes the $CH_2$ deformation band, showing that the 1401 $cm^{-1}$ (grey band) is genuinely a new peak in the Fh1-KO spectra, rather than an overall elevated Raman intensity.

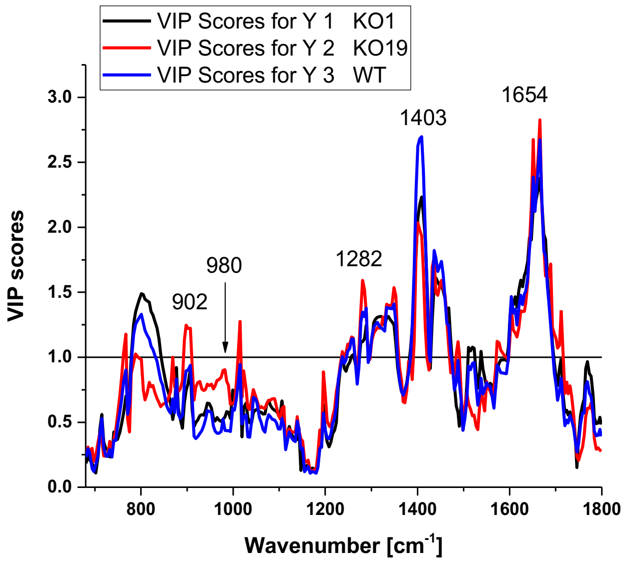


**Figure S9. PLS-DA can discriminate Fh1-WT and Fh1-KO cells using 532 nm excitation**. Graph shows the variable importance projection scores for the PLS-DA model, which was built on spectra acquired from line scans taken through the major axis of individual cells. These spectra were acquired with at 532 nm excitation (40 cell line scans, 20 steps per line, equating to 800 spectra per cell line at 5s acquisition time). 20 spectra were removed from the KO cl. 19 data set for low quality.

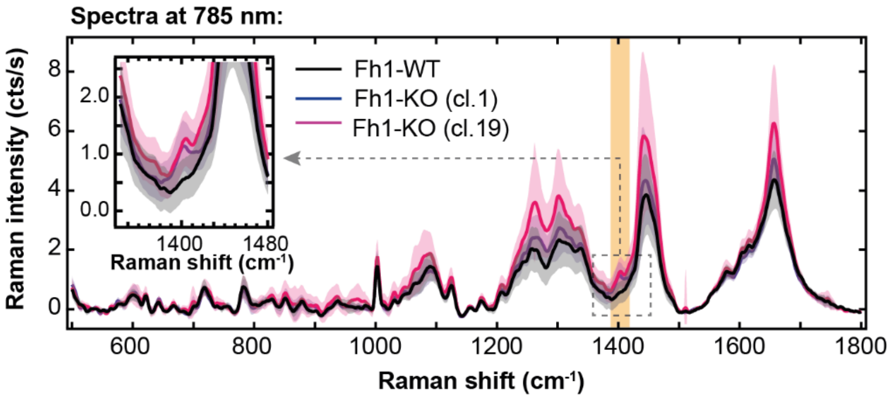


**Figure S10. Cell spectra acquired at 785 nm laser excitation.** Averaged line scan Raman spectra of Fh1-WT and Fh1-KO cells taken at excitation wavelength 785 nm (120 mW, 30x1s integration time, 10 steps per cell, 600 g/mm grating). Final spectra were smoothed with a Savitzky-Golay filter (7 point window) to reduce noise at this wavelength. Inset highlights the appearance of the 1401 $cm^{-1}$ peak in the Raman spectra of the two Fh1-KO clones.

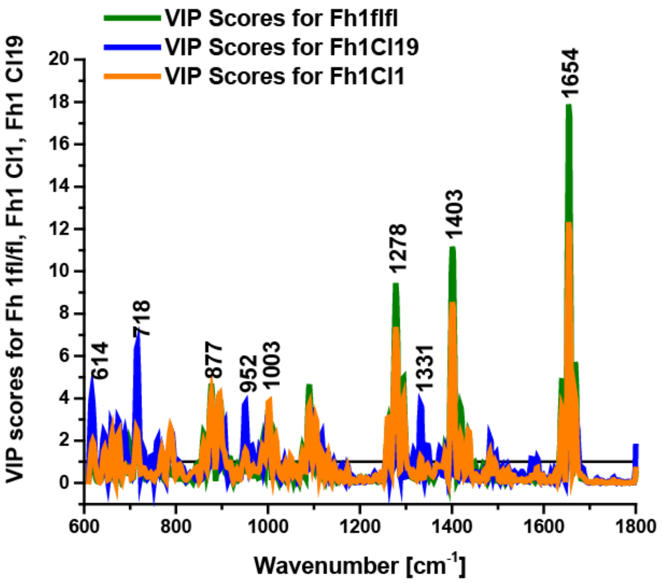


**Figure S11. PLS-DA can discriminate Fh1-WT and Fh1-KO cells using 785 nm excitation**. Graph shows the variable importance projection scores for the PLS-DA model, which was built on spectra acquired from line scans taken through the major axis of individual cells. These spectra were acquired with at 785 nm excitation (60 cell line scans, 10 steps per line, equating to 800 spectra per cell line at 30s acquisition time).

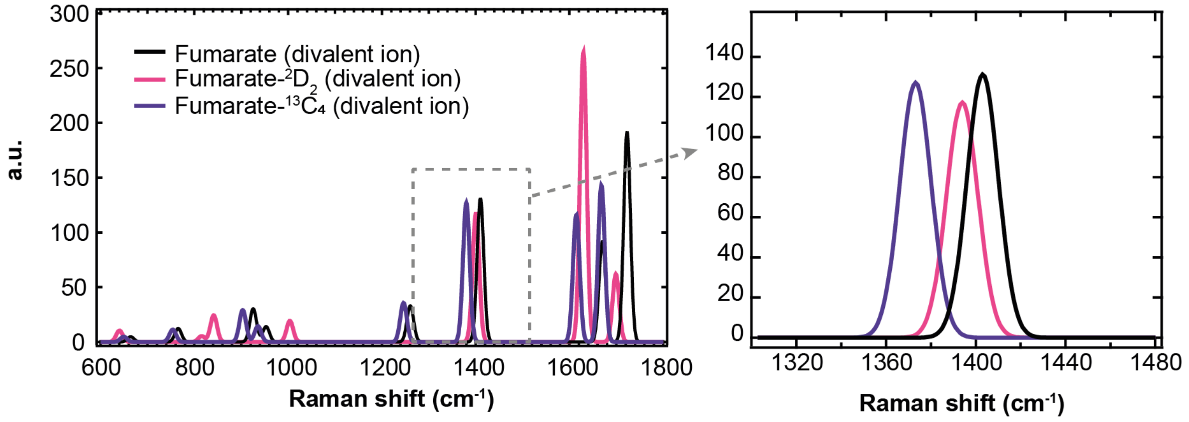


**Figure S12. Cells cultivated in isotopically labelled L-glutamine-$^{13}C_5$.** Raman spectra of isotopically labelled fumarate (fumarate-$^{13}C_4$) in fully ionized state as calculated by DFT.

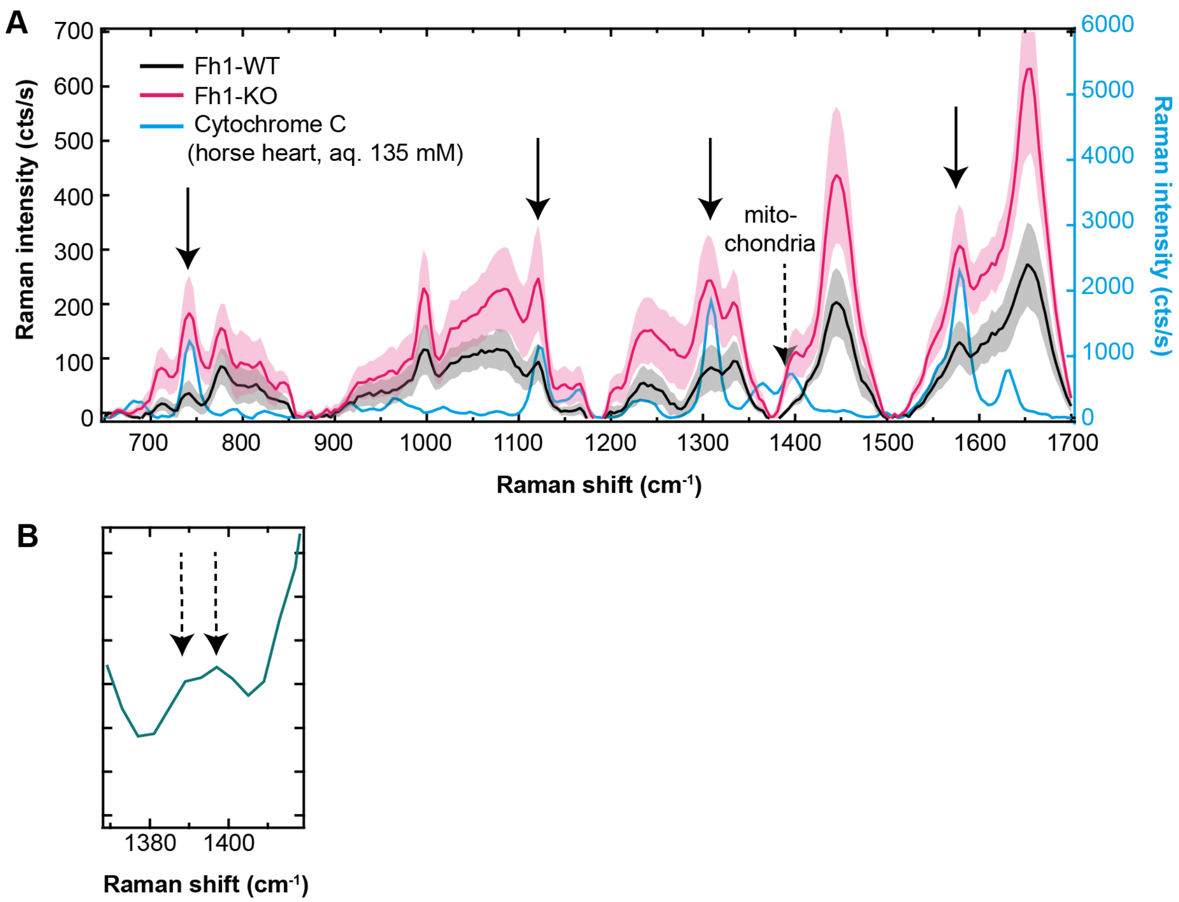


**Figure S13. A**) **Spectrum of Cytochrome C** (135 mM aqueous solution) overlayed on the averaged area scans for Fh1-WT and Fh1-KO cells, showing the bands related to cytochrome C (highlighted by solid arrows). The bands are strong on account of resonant Raman scattering for the incident wavelength $\lambda_{exc}$ = 532 nm. The dashed arrow points out a cytochrome C band which we believe is responsible for the 1389 $cm^{-1}$ band visible in the mitochondria. **B**) Zoom-in of the mitocondria class of Fh1-WT cells from Figure 3B, to show the bands related to oxidized and reduced Cytochrome C.

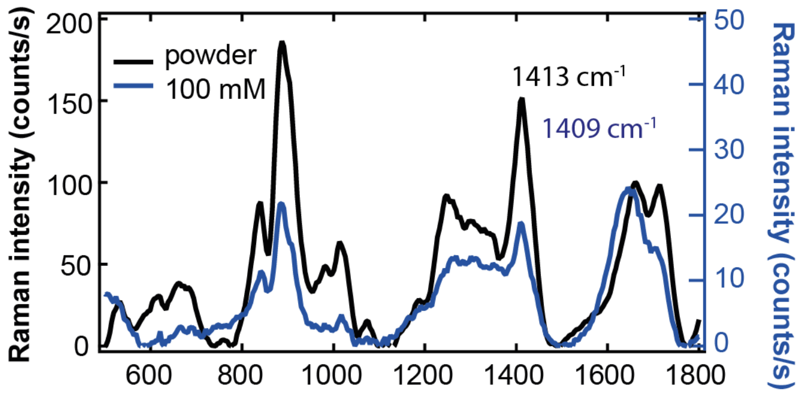

S-(1,2-Dicarboxyethyl)glutathione

**Figure S14. Raman spectra of S-(2-succinyl)glutathione** as powder (black trace) and a 100 mM aqueous solution (blue trace) at 532 nm, 30 mW laser illumination.

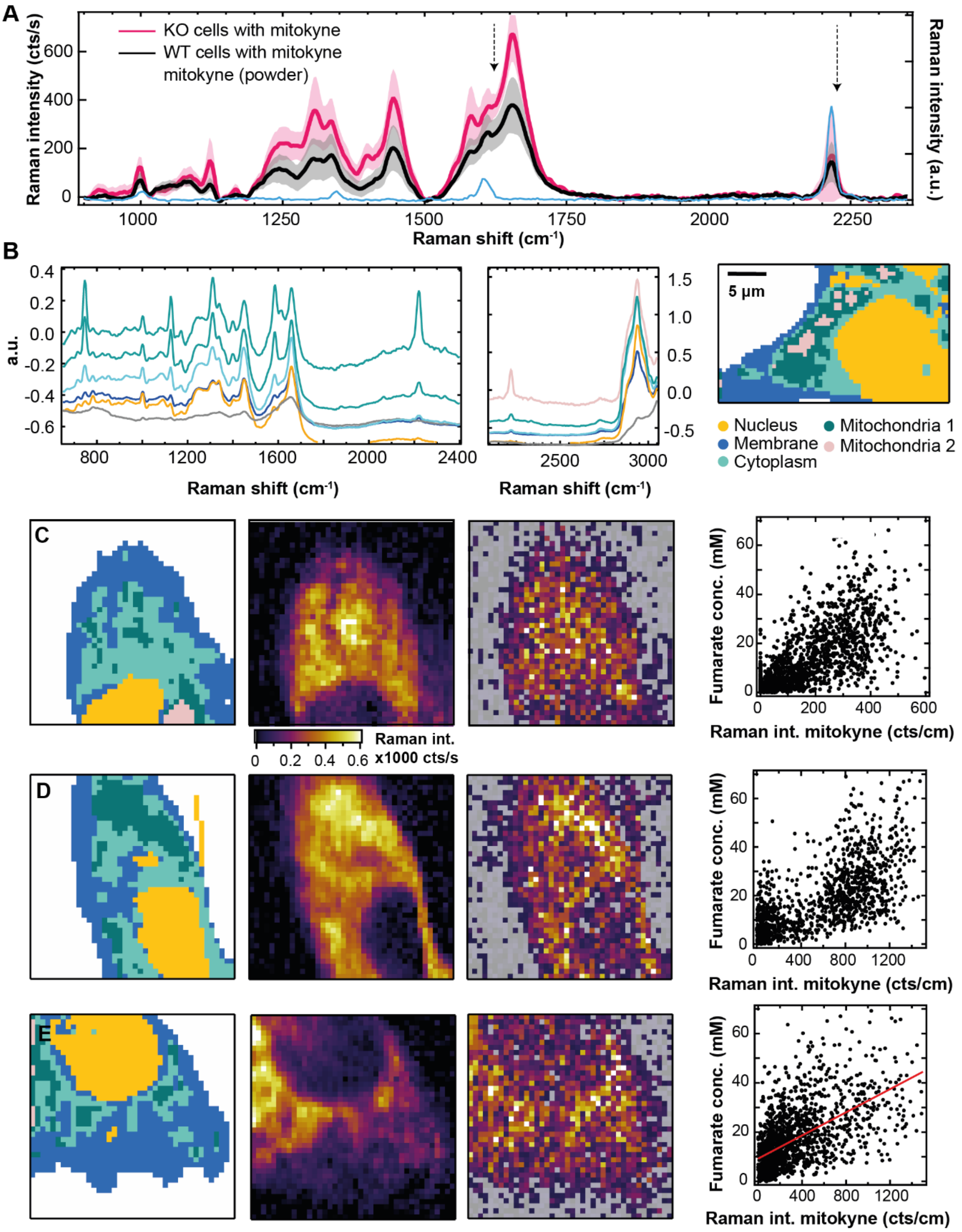


**Figure S15. Correlating mitochondrial assignment with k-means clustering using mitokyne.** (**A**) Average cell spectrum of Fh1-WT and Fh1-KO cells tagged with mitokyne, showing the primary and secondary band on top of the cell spectrum indicated by dashed arrows. (**B**) Loadings of a joint k-means clustering analysis over KO cells with (10 cells) and without (20 cells) mitokyne, showing that loadings corresponding to mitochondria have the highest mitokyne peaks. Two classes were assigned to the mitochondria. Cluster maps for non-labelled cells were identical to those found for k-means analysis in absence of mitokyne-labelled cells (c.f. Figure 3E). (**C-E**) K-means cluster maps (left), mitokyne peak intensity maps (second column), and apparent fumarate concentration maps (third column) for two Fh1-KO cells labelled with mitokyne, visually showing that highest intensities of the primary mitokyne peak coincide with the mitochondria and cytoplasm classes. Note that the mitokyne concentration is not calibrated, hence Raman intensities do not represent a mitokyne concentration. Occasional misassignment of cluster classes is evident, such as the two superfluous areas denoted as nucleus in (D). The fourth column shows scatter plots of the extracted fumarate concentrations versus mitokyne intensity, showing a positive correlation in each cell.

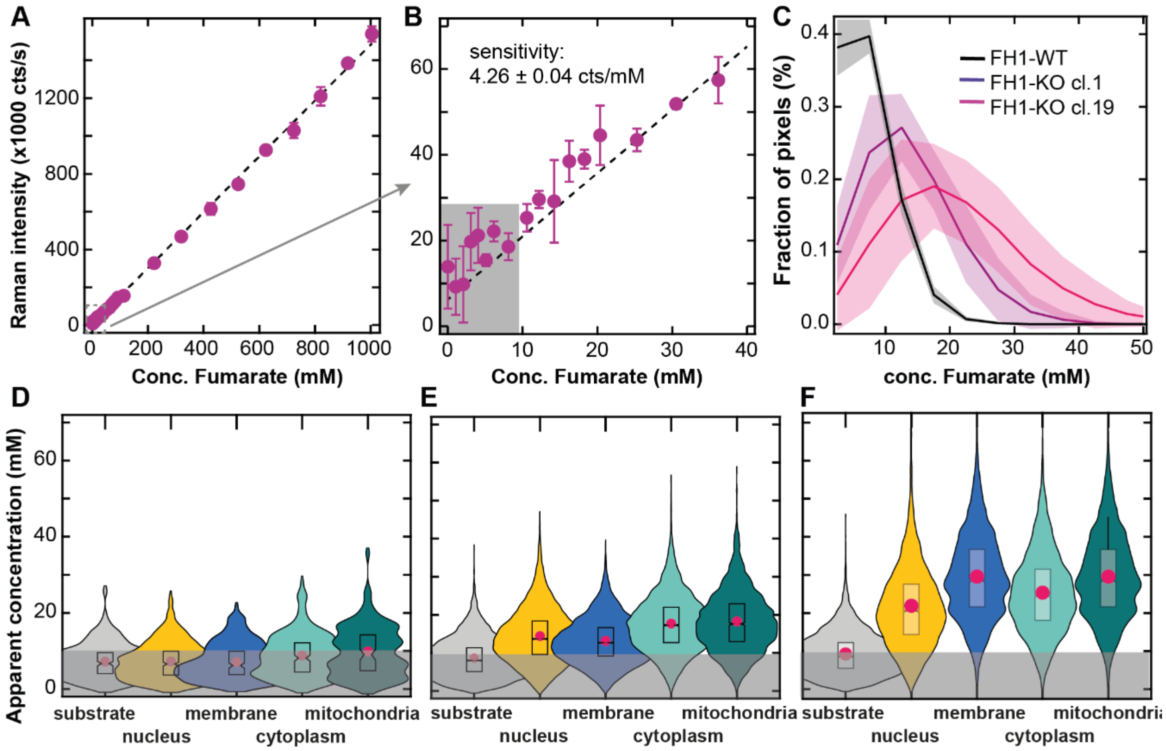


**Figure S16. Fumarate concentrations, determined by fitting a Gaussian peak shape** to the 1401 $cm^{-1}$ band (for $\lambda_{exc}$ = 532 nm, 26 mW power, 0.3 s dwell time). **A,B**) Raman intensity as a function of fumarate concentration for the 1401 $cm^{-1}$ peak. Error bars represent standard deviations of three separate measurements per concentration. r2 values of the linear regression is 0.998214. **C**) Fumarate concentrations found in n = 20 individual Fh1-WT cells (black) and n = 20 Fh1-KO cells (purple: cl.1, pink: cl.19) by fitting a Gaussian peak shape, normalized by the number of pixels per area scan (bin width: 5mM). Shaded areas represent standard deviation for each histogram bin. **D-F**) Fumarate concentrations per cluster class for all 20 Fh1-KO cells cl. 1 cells (D), 20 Fh1-KO cells cl. 1 cells (E), and 20 Fh1-KO cells cl. 19 cells (F). The pink dot indicates the mean fumarate concentration in the cluster class, while the box plot indicates the median value with 95% confidence interval (sloped edges), $1^{st}/3^{rd}$ quartile (box edges), and $9^{th}/91^{st}$ percentile whiskers. Grey area denotes concentrations below the LOD.

### Supplementary Videos

**Supplementary Video 1:** Animation of the C-H deformation (δ) mode of fully dissociated fumarate in vacuum, generated by DFT calculation (1261 $cm^{-1}$ band; 1277 $cm^{-1}$ observed experimentally).

**Supplementary Video 2:** Animation of the symmetric $CO_2^-$ stretch (v) mode of fully dissociated fumarate in vacuum, generated by DFT calculation (1410 $cm^{-1}$ band; 1401 $cm^{-1}$ observed experimentally).

**Supplementary Video 3:** Animation of the C=C stretch/$CO_2^-$ symmetric bending mode of fully dissociated fumarate in vacuum, generated by DFT calculation (1723 $cm^{-1}$ band; 1652 $cm^{-1}$ observed experimentally).

**Supplementary Video 4:** Animation of the C=C stretch/$CO_2^-$ asymmetric bending mode of fully dissociated fumarate in vacuum, generated by DFT calculation (1670 $cm^{-1}$ band).

**Supplementary Tables**

Table S1: Confusion matrices for spectra taken with a 532 nm laser

| **Model** | Actual Class | | |
|---|---|---|---|
| | KO 1 | KO19 | WT |
| Predicted as KO1 | 574 | 5 | 0 |
| Predicted as KO19 | 33 | 539 | 9 |
| Predicted as WT | 8 | 20 | 591 |
| Predicted as Unassigned | 0 | 0 | 0 |

| **Prediction** | Actual Class | | |
|---|---|---|---|
| | KO 1 | KO19 | WT |
| Predicted as KO1 | 190 | 4 | 0 |
| Predicted as KO19 | 11 | 172 | 5 |
| Predicted as WT | 4 | 13 | 195 |
| Predicted as Unassigned | 0 | 0 | 0 |

| **CV** | Actual Class | | |
|---|---|---|---|
| | KO 1 | KO19 | WT |
| Predicted as KO1 | 573 | 5 | 0 |
| Predicted as KO19 | 33 | 535 | 12 |
| Predicted as WT | 9 | 24 | 588 |
| Predicted as Unassigned | 0 | 0 | 0 |

| | KO 1 | KO19 | WT |
|---|---|---|---|
| Sensitivity (CV): | 0.925 | 0.936 | 0.973 |
| Specificity (CV): | 0.982 | 0.927 | 0.969 |

**Table 2:** Apparent average (mean ± standard deviation) fumarate concentrations for each cluster class, per cell line for Gaussian peak fitting.

| | Fh1-WT conc. (mM) | Fh1-KO Cl. 1 conc. (mM) | Fh1-KO Cl. 19 conc. (mM) |
|---|---|---|---|
| Substrate | 7 ± 4 | 9 ± 5 | 9 ± 5 |
| Nucleus | 7 ± 5 | 14 ± 7 | 22 ± 12 |
| Cell membrane | 7 ± 5 | 13 ± 6 | 30 ± 11 |
| Cytoplasm | 9 ± 6 | 18 ± 7 | 25 ± 11 |
| Mitochondria | 10 ± 6 | 18 ± 8 | 30 ± 11 |
