## Supplementary figures and images for "Raman micro-spectroscopy reveals the spatial distribution of fumarate in cells and tissues"

### Frequency 17_1261cm

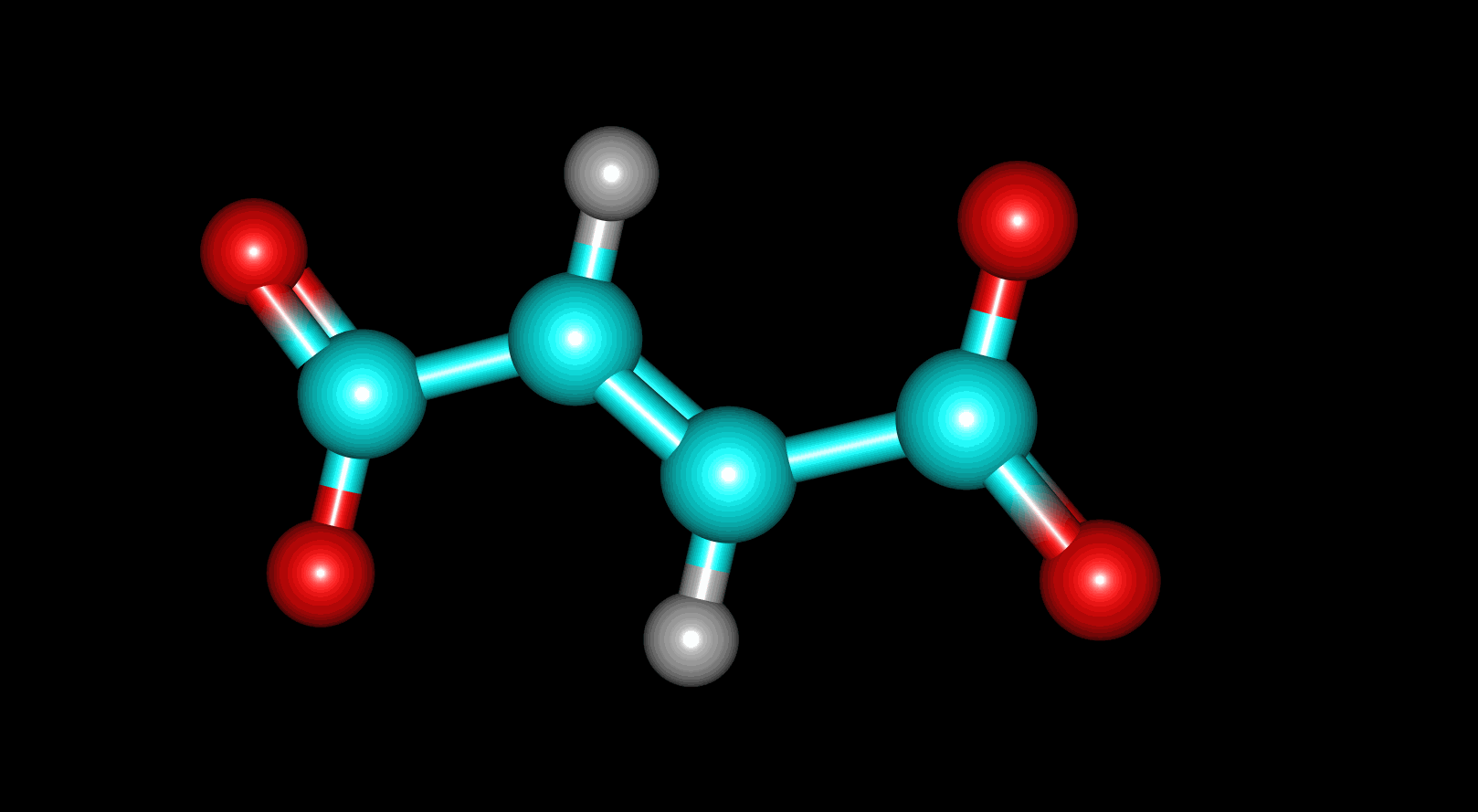

### Frequency 19_1410cm

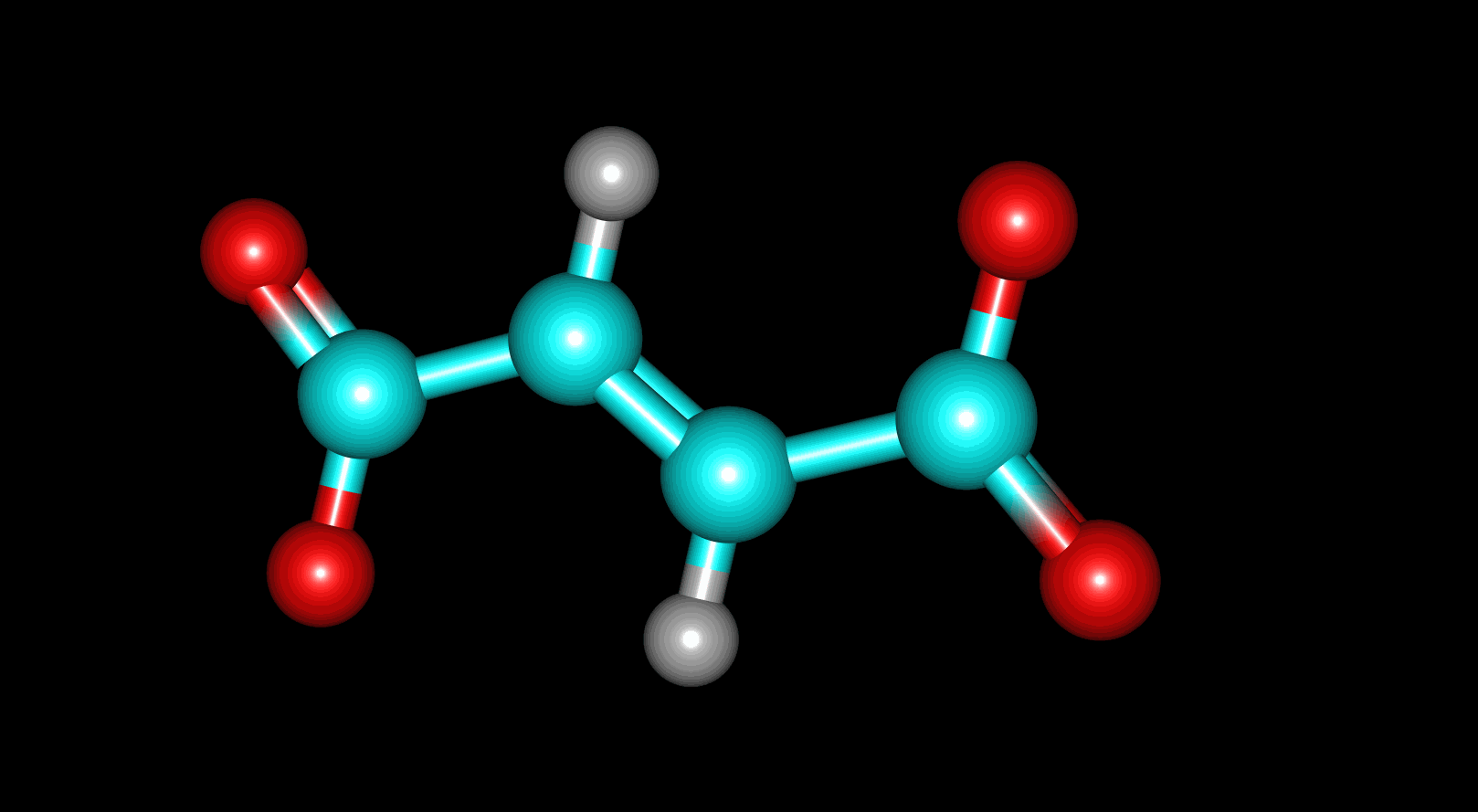

### Frequency 20_1670cm

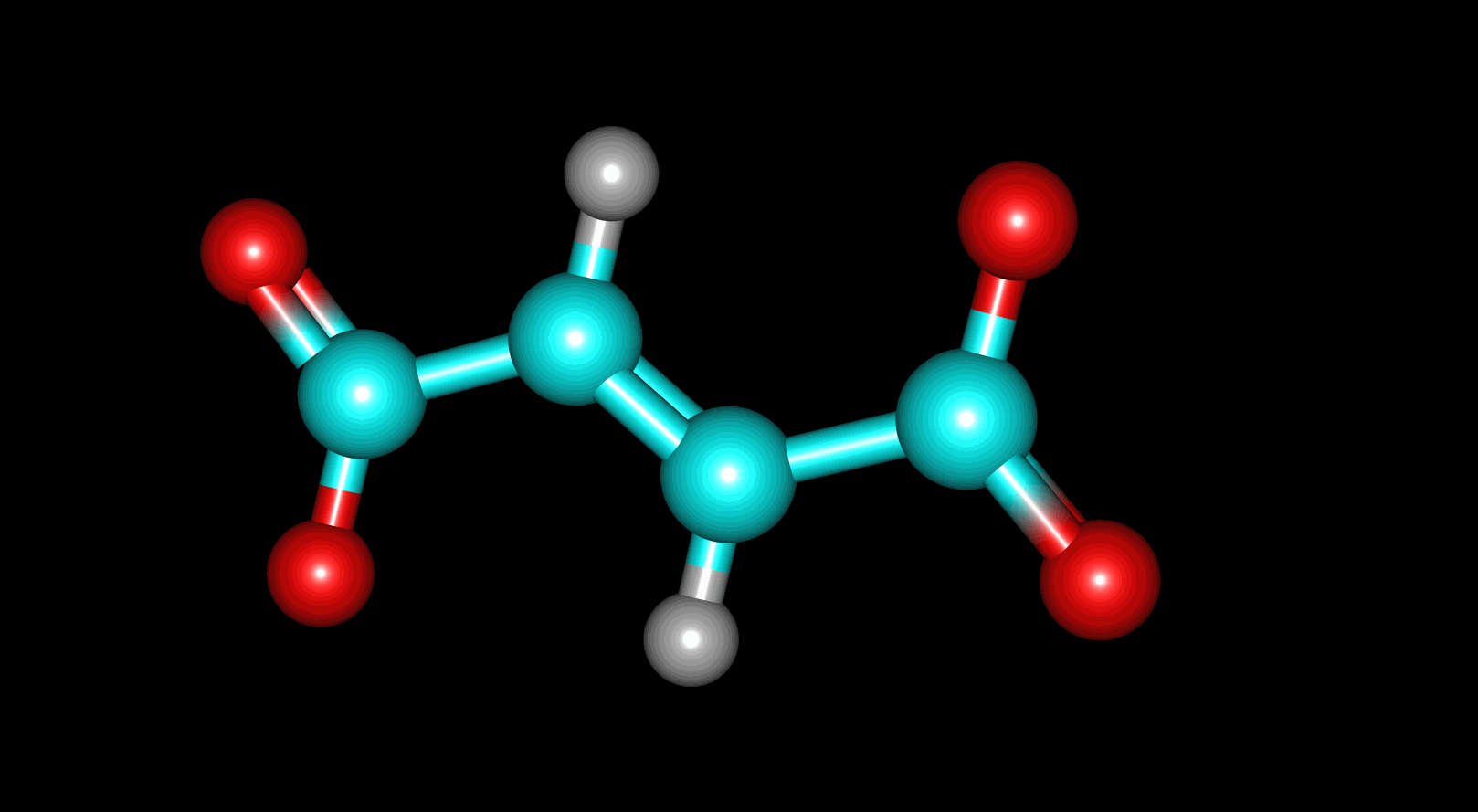

### Frequency 22_1723cm

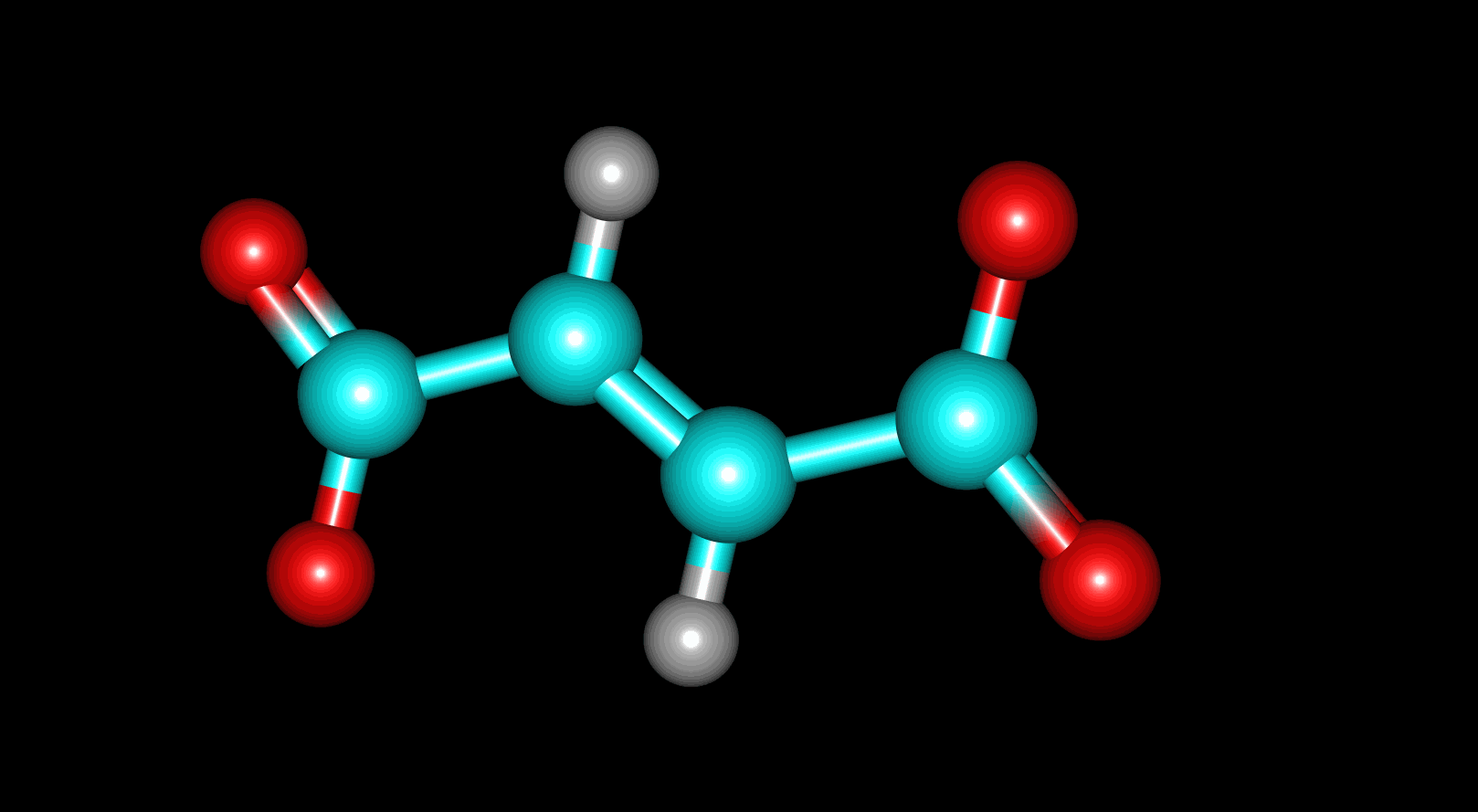
